# Detection of acute TLR-7 agonist-induced hemorrhagic myocarditis in mice by refined multi-parametric quantitative cardiac MRI

**DOI:** 10.1101/502237

**Authors:** Nicoleta Baxan, Angelos Papanikolaou, Isabelle Salles-Crawley, Amrit Lota, Rasheda Chowdhury, Olivier Dubois, Jane Branca, Muneer G. Hasham, Nadia Rosenthal, Sanjay K. Prasad, Lan Zhao, Sian E. Harding, Susanne Sattler

## Abstract

**BACKGROUND:** Hemorrhagic myocarditis is a potentially fatal complication of excessive levels of systemic inflammation. It has been reported in viral infection, but is also possible in systemic autoimmunity. Cardiac magnetic resonance (CMR) imaging is the current gold standard for non-invasive detection of suspected inflammatory damage to the heart and changes in T_1_ and T_2_ relaxation times are commonly used to detect edema associated with immune cell infiltration and fibrosis. These measurements also form the basis of the Lake Louise Criteria, which define a framework for the CMR-based diagnosis of myocarditis. However, they do not take into account the possibility of hemorrhage leading to tissue iron deposition which strongly influences T_1_ and T_2_ measurements and may complicate interpretation based on these two parameters only.

**METHODS:** Systemic inflammation was induced in CFN mice by application of the TLR-7 agonist Resiquimod. Histopathology was performed on heart sections to assess immune cell infiltration (Hematoxylin & Eosin), fibrosis (PicoSirius Red) and tissue iron deposition (Perl’s Prussian Blue). A multi-parametric cardiac MRI tissue mapping approach measuring T_1_, T_2,_ and T_2_^*^ relaxation times was established to non-invasively identify these parameters in this small rodent model.

**RESULTS:** Resiquimod-treated mice developed severe thrombocytopenia and hemorrhagic myocarditis. We identified patches of cardiac hemorrhage based on the presence of two major MRI phenotypes. Increased T_2_ with normal T_1_ and T_2_^*^ values correlated with infiltration/edema only, while decreased T_1_, T_2_, and T_2_^*^ values identify areas with infiltration/edema in the presence of iron indicating hemorrhagic myocarditis.

**CONCLUSION:** We show that over-activation of the TLR-7 pathway by Resiquimod treatment of CFN mice induces an early immune response reminiscent of excessive systemic inflammation due to viral infection. This causes internal bleeding which manifests most prominently as severe hemorrhagic myocarditis. We optimized a comprehensive noninvasive *in vivo* MRI approach based on quantitative measurement of T_1_, T_2_ and T_2_^*^ to demonstrate the presence of diffuse myocardial edema, infiltration and iron deposition without the need of contrast agent administration. We propose that adding quantitative T_2_^*^ mapping to CMR protocols for detection of myocarditis will improve diagnostic sensitivity and interpretation of disease mechanisms.

## 1) Introduction

Immune-mediated damage to the heart may occur as the result of a variety of underlying conditions including infectious disease, exposure to toxins, chemotherapeutic agents, immune checkpoint inhibitors and systemic inflammation due to autoimmune diseases. Besides a range of viruses known to cause severe hemorrhagic fevers, viral myocarditis can also be a complication of more common viruses, including most prominently coxsackie virus (1), adenovirus (2) or influenza (3). In rare cases, these can trigger excessive systemic inflammation and bleeding, which may manifest as hemorrhage in internal tissues and organs, including the heart. Several incidences of cardiac hemorrhage due to viral myocarditis have been described. Patchy cardiac hemorrhage has been detected in autopsy cases after myocarditis associated with swine-origin influenza (4)(5). Acute hemorrhagic pericarditis due to *Chlamydophila pneumoniae* was diagnosed in a pediatric pneumonia patient (6) and post-mortem cases of hemorrhagic myocarditis due to adenovirus infection have been reported (7). Acute hemorrhagic myocarditis has also been identified in a patient with systemic lupus erythematosus (8).

In addition, despite clearance of the initial viral insult, inflammation may persist due to the development of self-directed immune responses leading to persistent inflammatory cardiomyopathy characterized by myocardial contractile dysfunction similar to dilated cardiomyopathy (9).

Although, clinically detected cardiac hemorrhage may be considered a rare occurrence in myocarditis or systemic inflammatory autoimmune conditions (8), inflammatory effects on hematological parameters and vasculature may cause subclinical vessel damage and red blood cell extravasation which may need to be considered for accurate diagnosis. Cardiac magnetic resonance (CMR) imaging allows for non-invasive *in vivo* tissue characterization and is used to directly inform on cardiac involvement in a range of diseases (10)(11).

In this study, we further characterized the phenotype of acute cardiac hemorrhage in a mouse model of the autoimmune disease systemic lupus erythematosus (SLE) induced by application of the TLR-7 agonist Resiquimod (12). We made use of CFN mice, which we recently described to be highly susceptible to acute early inflammatory damage and subsequent inflammatory cardiomyopathy (13). We established a protocol of contrast agent free CMR parametric mapping for the detection of diffuse immune-mediated damage, taking into account a potential presence of hemorrhage.

## 2) Materials and Methods

### Mice and *in vivo* treatment

All mouse procedures were approved by the Imperial College Governance Board for Animal Research and in accordance with the UK Home Office Animals (Scientific Procedures) Act 1986 and Directive 2010/63/EU of the European Parliament on the protection of animals used for scientific purposes. Experimental mice were 10-12 week old littermates. They were housed in individually ventilated cages in temperature-controlled facilities on a 12 hour light/dark cycle on standard diet. The mouse line used in these studies was derived from a triple cross between C57BL/6J, FVB/NJ and NOD/ShiLtJ parental lines (CFN line) and displays high sensitivity to TLR-7 agonist Resiquimod (R848, Sigma-Aldrich, Dorset, UK) treatment (13). Treatment with Resiquimod was performed by topical application (100 μg/30 μl per 30 g body weight in 1:3 ethanol:acetone) to the ear three times a week as previously described (13).

### CMR

Mice were anesthetized and maintained under inhalation anaesthesia via a nose cone (2% isoflurane/medical oxygen). Respiration, ECG and body temperature were continuously monitored (1030-MR, SA Instruments, Stony Brook, NY, USA) through the CMR scans. Three ECG leads (SA Instruments, Stony Brook, NY) were placed subcutaneously on the left and right side of the thorax and on the right back leg and animals were positioned prone in a dedicated mouse bed. Body temperature was maintained between 36.5° and 37° by a circulating warm water heat mat.

All CMR scans were performed on a pre-clinical 9.4 T scanner (94/20 USR Bruker BioSpec; Bruker Biospin, Ettlingen, Germany) housed at the Biological Imaging Centre, Imperial College London using an 86 mm inner diameter volume transmit quadrature coil combined with an actively decoupled mouse heart array receiver. Data were acquired with Paravision 6.0.1 (Bruker, BioSpin).

For localisation of the heart, low-resolution ECG and respiratory triggered gradient echo scout scans were acquired in axial, sagittal and coronal orientations followed by pseudo two-and four-chamber views. This allowed reproducible planning of imaging slices in the true short axis orientation at three specific locations in the left ventricle: basal, mid and apical. A multi-parametric MR imaging approach was adopted and included mapping of T_1_, T_2_ and T_2_^*^ relaxation times to identify and characterize quantitatively the markers of inflammatory tissue injury observed in the Resiquimod-treated mice. For welfare reasons, due to the high susceptibility of Resiquimod-treated mice to acute bleeding, no contrast agent-based protocols were performed in this study.

*T_1_ mapping*: T_1_ mapping was performed (day0: n=12; 2.5w: n=9) using a gradient echo-based look-locker inversion recovery sequence with 20 inversion times (TI) which followed an adiabatic global inversion time. All inversions were R wave triggered to allow images to be acquired at the same part of the cardiac cycle (end diastole), with the TI points restricted to multiples of the R-R interval (RR ∼ 100 ms). Additional acquisition parameters were: inversion repetition time: 5 s (to allow full relaxation between inversions), TR = 4.5 ms, TE = 2.1 ms, flip angle 7°, slice thickness 1 mm, field of view = (22 × 22) mm^2^, spatial resolution (164×164) μm^2^, with 3 slices acquired sequentially in ∼ 14 min. The inversion points acquired during respiratory motion were manually removed. T_1_ curve fitting was subsequently performed on a pixel-wise basis on the remaining measured inversion times using a non-linear least square three-parameter fit which accounted for Look-Locker correction. Data analysis was performed in *Segment version 2.0* (Segment, Medviso) (14).

*T_2_ mapping:* T_2_ mapping was performed (day0: n=10; 2.5w: n=10) using a multi echo spin-echo fat suppressed sequence with 5 echo times acquired at the same part of the cardiac cycle (proximal to end systole). A pair of flow saturation bands surrounding each imaging slice were placed upstream the blood flow to minimize misleading signal dropouts caused by flow artefacts. Additional acquisition parameters were: TR/TE = 3500/2.74, 5.49, 8.23, 10.98, 13.72 ms, echo spacing = 2.74 ms, slice thickness 1 mm, field of view = (17 × 19) mm^2^, spatial resolution (147×167) μm^2^, GRAPPA acceleration factor 1.65, with 3 slices acquired sequentially in a total scan time of ∼ 12 min. A 2-parameter pixel-wise T_2_ fit was done assuming a mono-exponential signal decay. Myocardial segmentation excluded the regions of bright signal caused by the stagnant sub-endocardial blood occasionally observed on the T_2_ maps (15). Data analysis was performed in *Segment 2.0* (Segment, Medviso).

*T_2_^*^ mapping:* A multi-echo double cardiac and respiratory triggered 2D radial UTE sequence was used for T_2_^*^ mapping (day0: n=5; 2.5w: n=3). For this, the sequence was performed repeatedly with various TEs (0.37 ms, 1.5 ms, 2.2 ms, and 3 ms) in fixed scale of receiver gain. The other imaging parameters were as follows: TR = 5.6 ms, 12º flip angle, FOV = (20 × 20) mm^2^, in plane spatial resolution (164 × 164) µm^2^, 1 mm slice thickness, bandwidth 250 kHz, 352 number of projections, scan time of 3 min 30 s per each individual echo image. Precise k-space trajectory measurement is crucial to obtain good quality images when radial encoding is employed (16). Consequently, in this study, the radial trajectories were acquired in vivo from the signal of off-centered spins measured while playing out the gradient shapes in X, Y and Z directions, and measuring the phase difference of the retrieved signal (16). To ensure that the measured trajectory is identical to the actually used imaging gradients, the ADC delay and the ramp-up shape of the readout gradient were included in the trajectory calibration. To improve the accuracy of the *in vivo* trajectory measurement, we used 12 averages and ECG triggering. A pixel wise T_2_^*^ mono-exponential fit was performed on the multiple echo T_2_^*^ w-UTE images using *Segment 2.0* (Segment, Medviso) (17).

*CMR Data analysis:* Regional tissue characterization was performed on the T_1_, T_2_ and T_2_^*^ maps after partitioning the left ventricle wall based on the 16-segmentation model as per recommendations by the American Heart Association (six segments for basal and mid-levels, four segments for the apex) (18). Epicardial and endocardial contours were traced manually to define the myocardial wall of each animal (19). Particular care was taken to avoid signal contamination from the LV blood pool by excluding the innermost 5% of the myocardial wall. A layered pattern of signal dropout was observed surrounding primarily the sub-epicardial regions of the intra-ventricular septum in most of the mice. To account for this, each of the 6 cardiac segments of the mid-ventricular slice was divided further in two equal sections (sub-epicardial and sub-endocardial, average pixel number per segment of 36±5), resulting in 12 segments per slice (Fig. 2). Signals of each of the 12 segments were normally distributed, therefore T_1_, T_2_ and T_2_^*^ average values were computed and correlated with histology as described below. Correlation of MRI to histology data was performed using midline cross sections values. Voxel intensity histograms of the T_1_ and T_2_ maps were extracted over the full myocardial thickness of the mid ventricular anterior septum and anterior lateral walls of both Resiquimod-treated and untreated mice. Histograms of T_2_^*^ maps were afterwards compared to their respective T_1_ and T_2_ histograms to further evaluate their interdependence and to better characterize their specific distribution pattern in the presence of inflammatory tissue injury and myocardial bleeding. Histograms were reconstructed using 20 bins, mean or median, as appropriate, were measured and used for further analysis. To better visualize the specific features of signal distribution of each MR mapping technique, smoothing was performed by averaging 2 values on each side and using a second order polynomial smoothing (20).

### Macroscopic observations and scoring

Hearts were perfused and excised and macroscopically visible hemorrhagic lesions were scored on a scale from 0; no visible lesions, 1; lesions cover <10 % of heart surface, 2; lesions cover 10-30 % of heart surface, 3; lesions cover 30–50 % of heart surface, to 4; lesions cover >50 % of heart surface.

### Histology and scoring of damage parameters

Hearts of treated and untreated mice were excised after *in situ* perfusion with ice-cold Phosphate buffered saline (PBS, Sigma-Aldrich, Dorset, UK) through the apex of the left ventricle of the heart to clear blood from heart chambers and blood vessels and fixed in 4 % formaldehyde overnight, dehydrated in an increasing gradient of ethanol and embedded in paraffin. Five µm sections were cut and de-waxed and rehydrated in an ethanol gradient. Sections were stained with haematoxylin and eosin (H&E), PicoSirius Red and Perls Prussian Blue stain. All reagents were purchased from Sigma Aldrich (Sigma-Aldrich, Dorset, UK). Semi-quantitative scoring was performed as established previously (13). Hematoxilin & eosin stained sections were used to analyze extravasation of red blood cells and mononuclear cell infiltration. PicoSirius Red staining was used to analyze and score fibrosis. Perls Prussian Blue staining was used to analyze and score cardiac iron deposition. Individual parameters were scored on a scale of 0 (none), 1 (mild), 2 (moderate) to 3 (severe). Scores were obtained from 12 segments along the myocardium at a 100x magnification on four midline cross sections per animal by a blinded researcher. Images were captured using a LMD7000 microscope (Leica microsystems, Milton Keynes, UK) and processed using the public domain software ImageJ (NIH; http://rsb.info.nih.gov,) (21).

### Hematological analysis

Mouse blood was collected into 129 mM trisodium citrate and platelet counts were determined immediately by flow cytometry using a rat antibody recognising mouse platelet GPIbβ (Emfret Analytics, Eibelstadt, Germany) and calibrated beads (Saxon Europe, Kelso, UK) according to manufacturer’s instructions. Samples were analyzed using a FACScalibur flow cytometer. Hemoglobin content in blood samples was determined by the cyan-methemoglobin method using Drabkins reagent and bovine hemoglobin as a standard (both Sigma-Aldrich, Dorset, UK). Blood or hemoglobin standards were diluted in 200 µl Drabkin/Brij L23 solution, incubated at room temperature for 15 min and absorbance read at 540nm. Hemoglobin content in mouse blood samples were extrapolated from a standard curve of bovine hemoglobin by linear regression using GraphPad Prism (v7).

### Experimental design and statistical analysis

Animal number and sample sizes calculations were performed using G* Power 3.1 (22) available at http://www.gpower.hhu.de/ and reflect effect sizes obtained in previous experiments with comparable readouts. Scoring of histopathology was performed by a blinded researcher. Statistical analyzes were performed using SPSS or GraphPad Prism. Normally distributed data were presented as mean±s.e.m. Mann-Whitney test was used for scoring data, and one-or two-tailed unpaired or paired Student’s t-tests were performed as appropriate for parametric data.

## 3) Results

### 3.1) TLR-7 agonist Resiquimod induces severe pan-cardiac hemorrhage

As shown previously (13), Resiquimod-treatment induced patches of cardiac hemorrhage which were severe enough to be macroscopically visible on the surface of the heart in the majority of treated CFN mice (Figure 1A, B). Histopathological examination reveals immune cell infiltration, edema, cardiomyocyte damage and accumulation of red blood cells (RBC) between myocardial fibers (Figure 1C). Microscopically, RBC extravasation into the surrounding tissue is evident in all treated mice (Figure 1C, graph). Resiquimod-treated mice also develop clinical hematological manifestations including thrombocytopenia and anemia (Figure 1E) after only two weeks of treatment.

**Figure 1:**
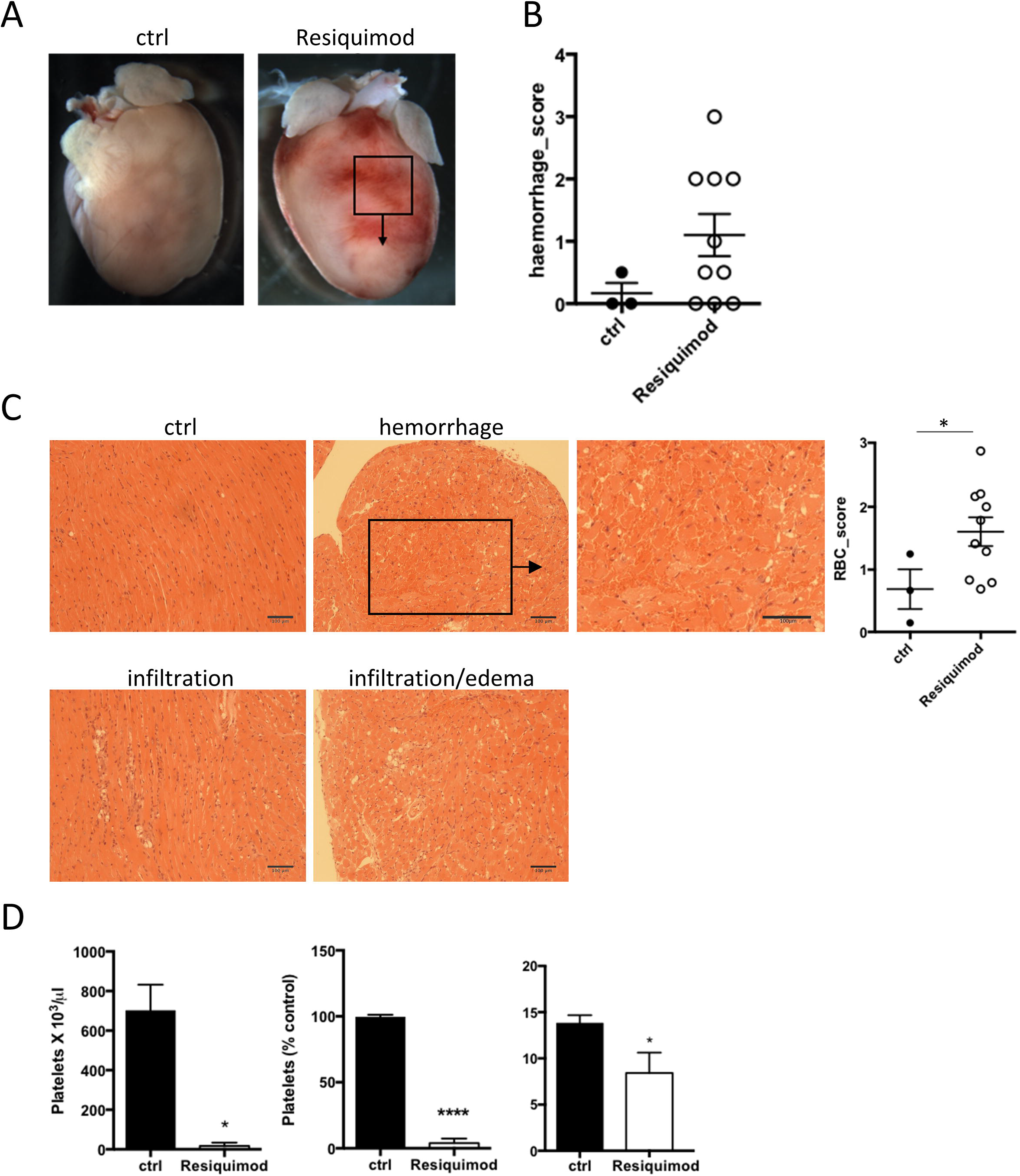
TLR-7 agonist Resiquimod induces myocarditis, thrombocytopenia and cardiac hemorrhage. Mice were treated with Resiquimod by topical application to the ear three times a week for two weeks. Cardiac involvement was assessed macroscopically and by microscopic histopathology. A: Example of macroscopic hemorrhagic lesions on hearts of Resiquimod-treated mice. B: Hemorrhagic lesions were scored on a scale according to: 0: no visible lesions, 1 lesions cover <10 % of heart surface, 2 lesions cover 10-30 % of heart surface, 3 lesions cover 30–50 % of heart surface, 4 lesions cover >50 % of heart surface. C: Micrographs of Hematoxylin & Eosin stained paraffin-embedded heart sections of Resiquimod-treated mice showing histopathological damage parameters including immune cell infiltration and associated edema as well as red blood cells in the myocardial interstitial space. The degree of RBC extravasation per animal was assigned a global semi-quantitative score of 0 to 3. D: Hematological analysis was performed to obtain full platelet count, percentage drop in platelet number from baseline in response to Resiquimod-treatment and changes in hemoglobin content in response to Resiquimod-treatment. Mann-Whitney was used for semi-quantitative histopathology scores; n=3 (ctrl), 10 (treated), unpaired Student’s t-test for hematological values; n=3, * P<0.05, * * P<0.005 and * * * P<0.001.

### 3.2) Native CMR T_1_ and T_2_ tissue mapping does not conclusively correlate with inflammatory cardiac damage in Resiquimod-treated mice

Resiquimod-induced systemic inflammation leads to myocarditis as assessed by histopathology (Figure 2A). We therefore initially set out to measure cardiac inflammatory damage parameters (infiltration/edema and fibrosis) by classical CMR T_1_ and T_2_ relaxation times. Global T_1_ and T_2_ values obtained by CMR imaging were extracted from the mid slice for comparison with histopathological changes in corresponding histology sections (Figure 2B). Global histopathological damage scores showed mild changes compared to baseline values for inflammatory infiltration and negligible changes in collagen deposition were observed in Resiquimod-treated compared to untreated animals due to the acute stage of disease (Figure 2C). Global T_1_ values show a significant decrease (T_1_ = 1128 ± 121 ms, p=0.017) compared to myocardial tissue of untreated mice (T_1_=1505 ± 18.5 ms) whereas T_2_ did not significantly change compared to controls (28.5 ± 0.5 ms vs 27 ± 3 ms) (Figure 2D).

**Figure 2:**
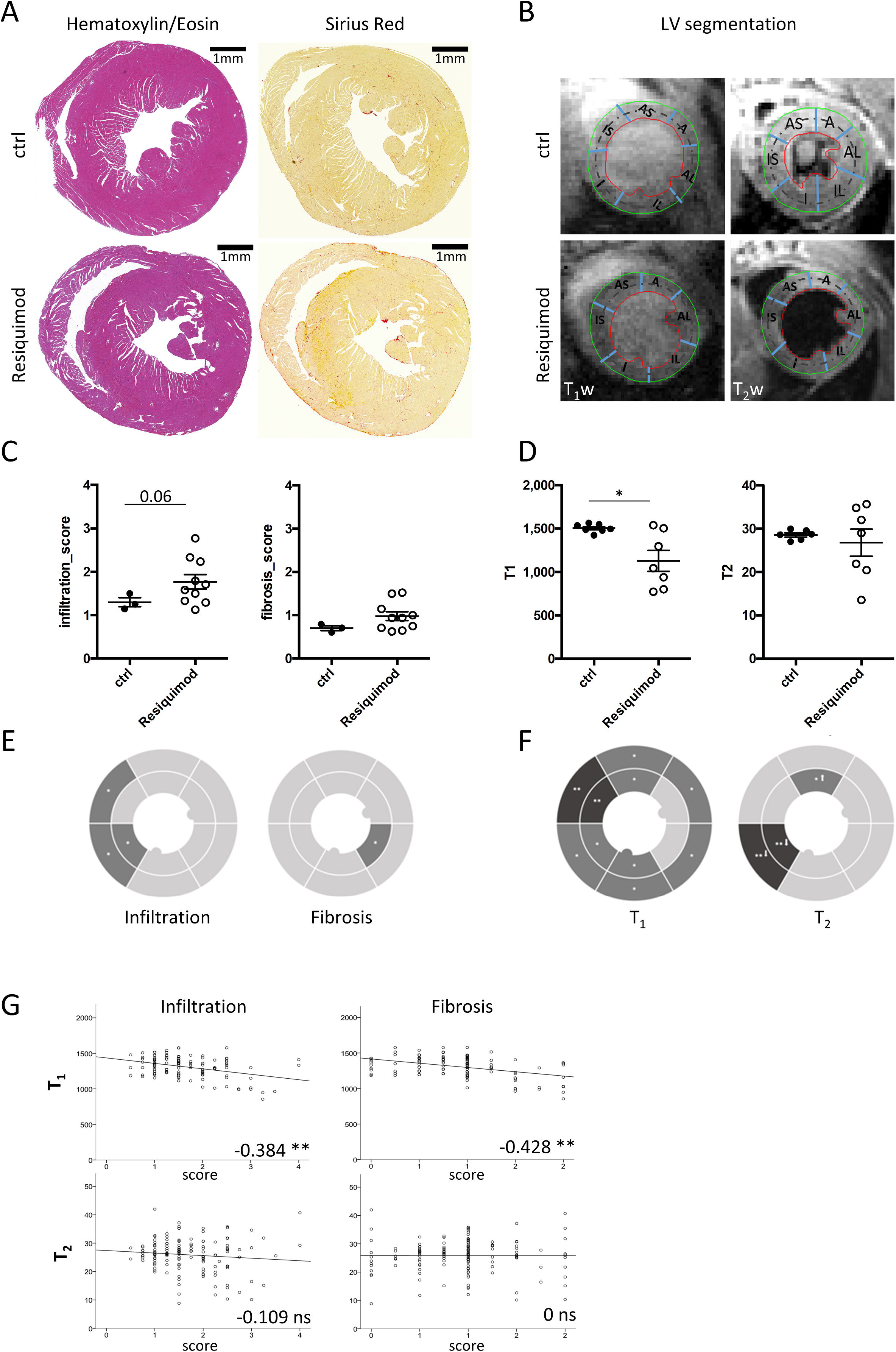
Native T_1_ and T_2_ tissue mapping does not conclusively correlate with inflammatory cardiac damage in Resiquimod-treated mice. Mice were treated with Resiquimod by topical application to the ear three times a week for two weeks and heart tissue damage was assessed by histology and MRI. A: Example of Hematoxylin & Eosin and PicoSirius Red stained paraffin-embedded heart sections of Resiquimod-treated and control mice showing significant immune cell infiltration and mild fibrosis. B: Example of corresponding T_1_w and T_2_w images to demonstrate segmentation of mid-section LV as per AHA recommendations: (A: anterior, AL: anterior lateral, IL: inferior lateral, I: inferior, IS: inferior septum, AS: anterior septum). C: Hematoxylin & Eosin and PicoSirius Red stained heart sections were given a global semi-quantitative score for infiltration, fibrosis and extravasation of red blood cells on a scale from 0-3 (none, mild, moderate, severe). D: Global values for T_1_ and T_2_ values of R848 mice compared to healty controls. E/F: Regional analysis depicted as pie charts performed on 12 segments at mid-level per mouse of infiltration, fibrosis, RBC accumulation (E), T_1_ and T_2_ (F). G: Correlation between T_1_ and T_2_ indices with individual histopathology scores. Statistics: Symbols in graphs represent the average score of individual mice, error bars show mean+/-s.e.m. Asterisks in pie charts depict significance of difference between Resiquimod-treated and control mice in the respective segment of the heart. Mann-Whitney was used for semi-quantitative histopathology scores; n=3(ctrl), 10(treated), paired Student’s t-test for longitudinal MRI values; n=7, * P<0.05, * * P<0.005 and * * * P<0.001. Pearson’s Correlation was used to assess significance of correlation pooling data from three independent experiments n=10-12 (ctrl), 9-10 (treated), * Correlation is significant at the 0.01 level (2-tailed). * * Correlation is significant at the 0.05 level (2-tailed).

These findings suggest that average values over the mid slice reflect an average between damaged and healthy areas, which may reduce sensitivity of detection. Severity of damage was most evident when regional analysis was performed on each of the 12 segments selected as described above (Figure 2E). Histopathological findings showed that the intra-ventricular septum and sub-epicardial areas are most severely affected, while LV free wall and sub-endocardium show mild damage only. Similarly, CMR imaging detected a significant drop of sub-epicardial T_1_ values in the Resiquimod-treated mice particularly in the anterior septum. Regional T_2_ values showed no significant change, except for the inferior septum (Figure 2F) were values decreased. Despite a seemingly similar spatial distribution pattern between changes in regional T_1_ and T_2_ values and corresponding histopathological scores (Figure 2E, 2F), pairwise Pearson’s correlation between CMR values and histopathology was mild but significant for T_1_ (infiltration: -0.384, fibrosis: -0.428) and not significant for T_2_ (infiltration: -0.109, fibrosis: 0) (Figure 2G, Table 1). For example, native T_1_ values decreased significantly in the intra-ventricular septum and along the entire sub-epicardium, which is contrary to the expected increase due to histologically observed immune cell infiltration (Figure 2E, 2F).

### 3.2) Regional T_1_ and T_2_ mapping is strongly influenced by iron deposition in the tissue

The finding that regional T_1_ and T_2_ values did not correspond well with histopathological scores prompted us to further assess their relationship. In the presence of edema/infiltration water content within the tissue is increased and T_2_ is expected to increase. Indeed, we identified areas of high T_2_ values (bright contrast in T_2_ map) which corresponded to areas of inflammatory infiltration and edema in H&E histology (Figure 3A, box A2). Yet, other areas with comparable levels of immune cell infiltration as per histology, showed unchanged or even decreased T_2_ values (Figure 3A, box A1). Fibrosis, has been suggested as another parameter to increase T_2_ values (23), however as levels of fibrosis were negligible and evenly distributed across the hearts due to disease stage, fibrosis was rejected as likely factor to cause differences in T_2_ values across the heart.

**Figure 3:**
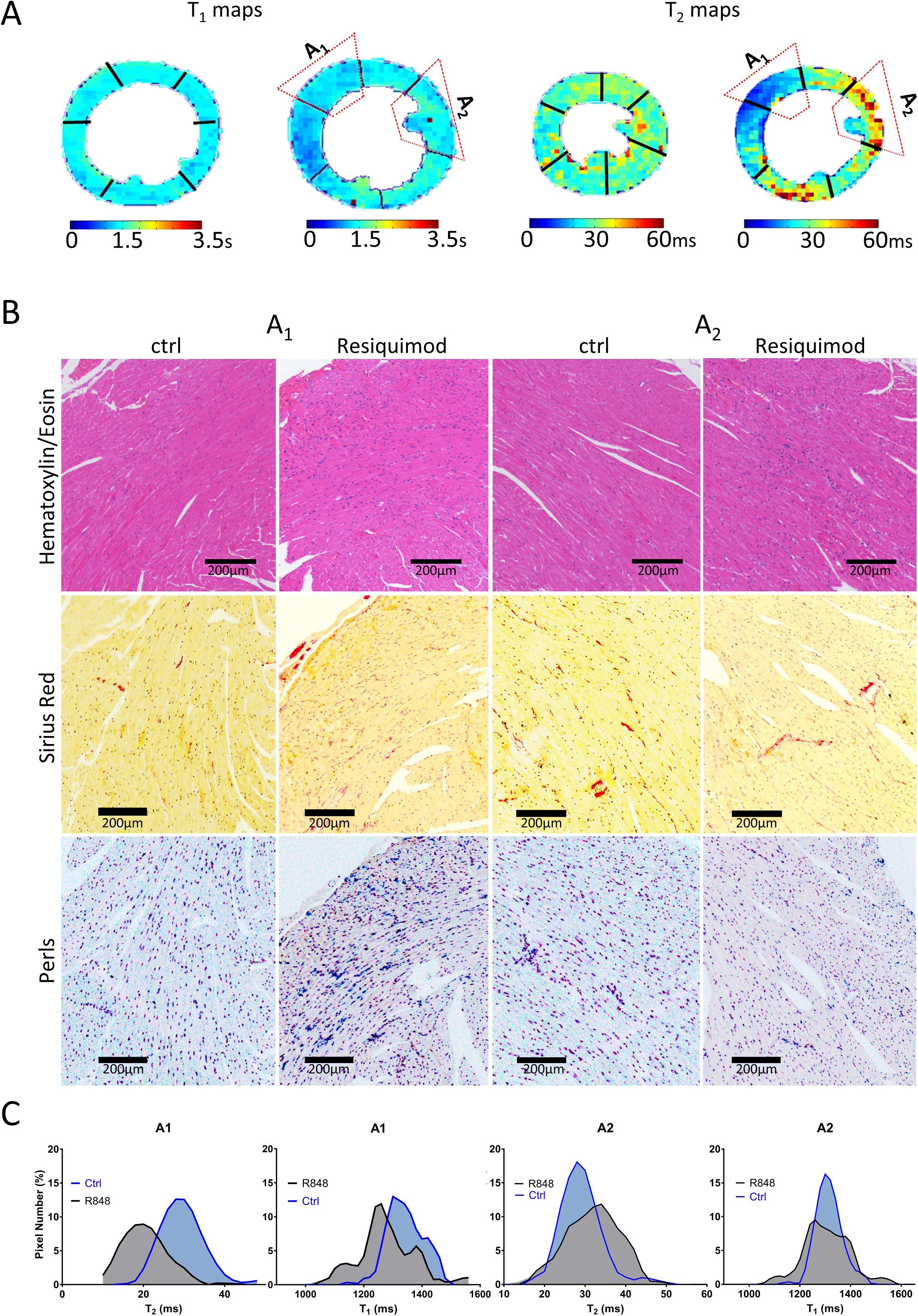
Regional T_1_ and T_2_ mapping is strongly influenced by iron deposition in the tissue. Mice were treated as described above and heart tissue damage was assessed by histology and MRI. A: T_1_ and T_2_ maps illustrating heterogeneity of T_1_ and T_2_ relaxation times with regions of low T_1_, T_2_ (A1) and high T_2_ (A2). B: Corresponding areas of interest (A1 and 2) in Hematoxilyn & Eosin, PicoSirius Red and Perls Prussian Blue stained paraffin-embedded heart sections. C: Histograms of signal distribution of T_1_ and T_2_ indices from A1 and A2 regions of interest.

The second dominant histopathological phenotype in hearts of Resiquimod-treated CFN mice, is the substantial accumulation of interstitial erythrocytes, which lead us to investigate iron deposition in the hearts. There was indeed a significant level of iron deposition in areas of particularly low T_1_ and T_2_ values (Figure 3B, box A1), while areas with less iron yielded increased T_2_ values in the presence of inflammation (Figure 3B, box A2).

Histograms of T_1_ and T_2_ signal distribution were computed from the anterior septum (A1) and anterior lateral (A2) wall (Figure 3C). The signal distribution of T_1_ mapping indices enabled a clear discrimination between regions of healthy myocardium and areas with high iron content (A1). T_2_ signal distribution behaved similarly confirming the dominant effect of iron in reducing T_1_ and T_2_ relaxation times.

In tissue areas with infiltration, but without iron deposition (A2), T_1_ showed high accuracy by not picking up the influences of T_2_ relaxation (24) and was characterized by a slightly more dispersed signal distribution compared to controls. T_2_ histograms presented a broader signal distribution with moderate increase of mean values compared to healthy control areas proving the sensitivity of T_2_ mapping to detect edema and cellular infiltrate only in areas without iron deposition.

In summary, iron deposition was the main reason for the observed decrease in T_1_ and T_2_ relaxation times. Tissue areas lacking iron deposits follow the classical paradigm that T_1_ detects fibrosis and T_2_ detects edema/cell infiltration.

### 3.3) An optimized T_2_^*^ mapping approach reveals iron deposition which impacts on T_1_ and T_2_ values

Due to its high sensitivity to paramagnetic iron, T_2_^*^ mapping has become the gold standard CMR measure to detect and quantify iron storage molecules (ferritin and hemosiderin) deposited in tissue (25). However, the LV free wall is usually excluded from T_2_^*^ measurements because of large magnetic susceptibility artefacts caused by the proximity to the air-filled lung. A remedy to reduce this susceptibility-induced fast T_2_^*^ decay is the application of an ultra-short echo time (UTE) readout instead of the conventional gradient-echo readout (26).

Using this approach, we obtained artefact-free T_2_^*^ maps with full coverage of the intra-ventricular septum and free wall at apical, mid and basal level of the mouse heart. An example of the high-quality T_1_, T_2_ and T_2_^*^ maps acquired in healthy controls is shown in Supplementary Figure 1.

RBC extravasation and iron accumulation increased in Resiquimod-treated mice, and global T_2_^*^ values dropped significantly (3.7 ± 0.2 ms, Figure 4A). Perls Prussian Blue staining showed a significant increase in tissue iron in myocardial regions with low T_2_ and T_2_^*^ relaxation times. Representative T_1_, T_2_ and T_2_^*^ maps of Resiquimod-treated mice with corresponding histological iron staining of heart sections are presented in figure 4B. Histology micrographs were magnified to show an example of severe iron accumulation in the anterior wall of the inter-ventricular septum. The corresponding PicoSirius Red-stained section demonstrate mild interstitial fibrosis and H&E stains show mononuclear cell infiltration (Figure 4C). Histograms of T_2_^*^ signal distribution reveal a skewed distribution towards lower median values consistent with the T_2_^*^ hypo-intense regions of strong iron deposition, whereas the histograms of T_1_ and T_2_ show low mean values but the normal distribution of signal suggest a weaker sensitivity to iron compared to T_2_^*^ (Figure 4D).

**Figure 4:**
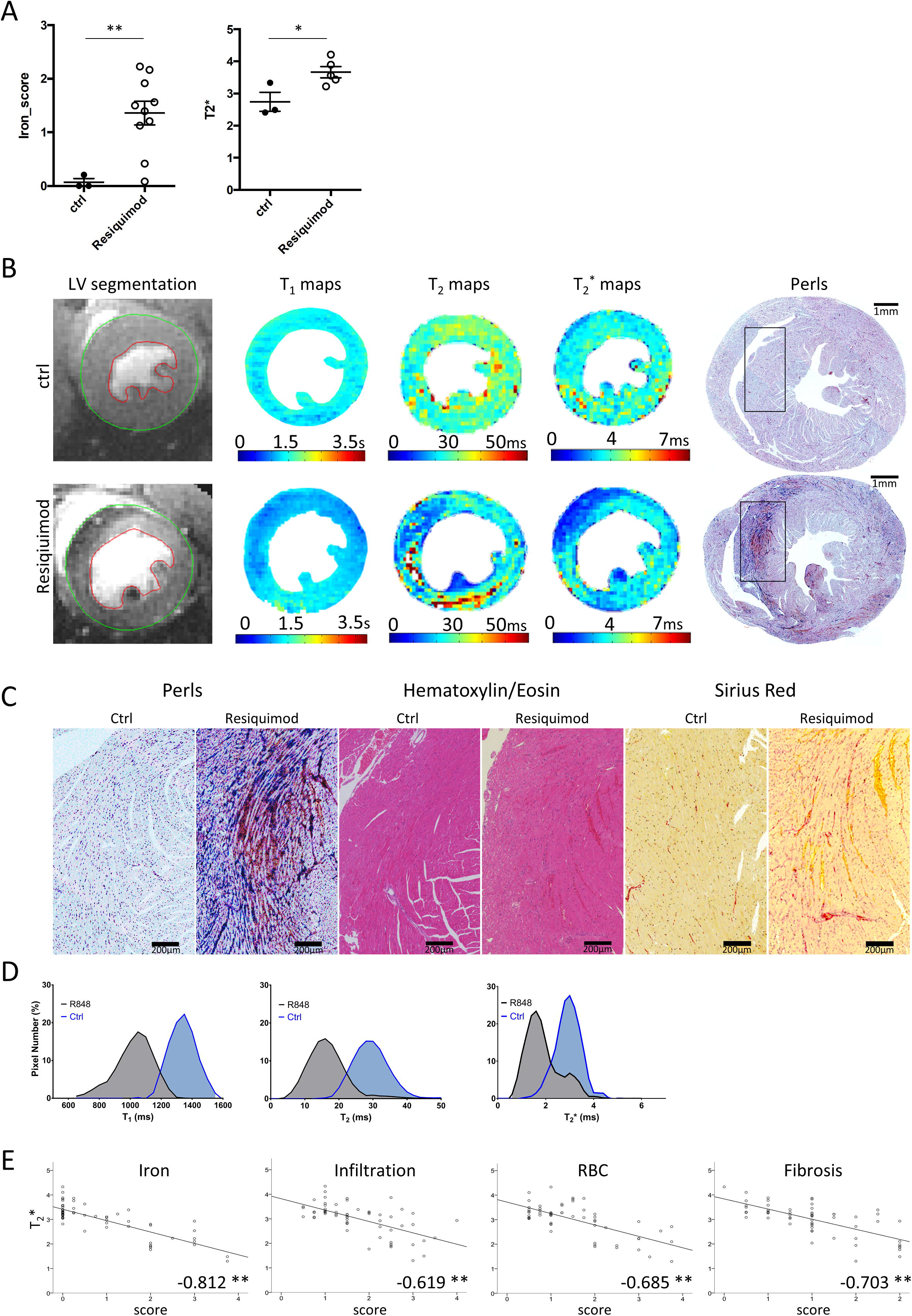
T_2_^*^ reveals severe cardiac iron deposition impacting on T_1_ and T_2_ measurements. Mice were treated as described above and heart tissue damage was assessed by histology and MRI. A. Global histopathological scores of RBC extravasation, Iron and T_2_^*^ values of Resiquimod-treated mice compared to control mice. Hematoxilyn & Eosin and Perls Prussian Blue stained heart sections were given a global semi-quantitative score for extravasation of red blood cells and iron deposition on a scale from 0-3 (none, mild, moderate, severe). B: Example of T_1_, T_2_ and T_2_^*^ maps of mid-section LV with corresponding Perls Prussian Blue-stained paraffin-embedded heart sections of Resiquimod-treated and control mice showing significant iron deposition. C: High magnification images of tissue areas with iron deposition (Perls Prussian Blue), immune cell infiltration (H&E) and mild fibrosis (PicoSirius Red). RBC infiltration scores are possible to obtain from each of these staining techniques. D: Histograms of T_1_, T_2_ and T_2_^*^ signal distribution in the anterior septum showing the impact of iron on the behavior of each relaxation time index. E: Correlation between T_2_^*^ values and individual histopathology scores. Statistics: Symbols in graphs represent the average score of individual mice, error bars show mean+/-s.e.m. Mann-Whitney was used for semi-quantitative histopathology scores; n=3(ctrl), 10(treated), unpaired Student’s t-test for MRI values; n=3 (ctrl), 5(treated), * P<0.05, * * P<0.005 and * * * P<0.001. Pearson’s Correlation was used to assess significance of correlation pooling data from three independent experiments n=3-5 (ctrl), 3-10 (treated), * Correlation is significant at the 0.01 level (2-tailed). * * Correlation is significant at the 0.05 level (2-tailed).

Correlation analysis between regional T_2_^*^ and histopathological parameters was performed and showed strong negative correlations with degree of iron deposition (-0.812), RBC extravasation (-0.685), infiltration (-0.619) and fibrosis (-0.703) (Figure 4E). Table 1 summarizes the correlation of all regional MRI indices to the respective histopathological findings. Most importantly, correlation between T_2_^*^ values and regional iron deposition scores is very strong and highly significant. T_1_ and T_2_ correlation with iron are -0.413 and -0.277, respectively. Correlation between T_2_^*^ and other damage parameters is lower and might be affected by a knock-on effect from their own correlation with iron deposition. Notably, neither T_1_, T_2_ or T_2_^*^ values changed in the liver of the same animals (Supplementary Figure 2), suggesting that inflammatory damage and hemorrhage are specific to the heart.

## 4) Discussion

Cardiac hemorrhage is, albeit considered rare, a potentially fatal complication of acute myocarditis (3). However, to date, a diagnosis is often only obtained post-mortem. Underlying mechanisms leading to hemorrhage in some myocarditis patients but not in others and implications on patient survival and subsequent development of inflammatory cardiomyopathies are not well understood. In addition, acute myocarditis may be a trigger for an aberrant autoimmune reaction against the heart (27) and it is feasible that severity of hemorrhaging, acute tissue damage and subsequent autoimmunity are correlated. In the brain, hemoglobin-derived iron plays a major role in secondary damage following hemorrhage (28) and iron-overload cardiomyopathy is caused by iron deposition in the tissue, induction of cardiomyocyte cell death and an ensuing inflammatory reaction (29). A recent study on experimental myocarditis in mice, detects iron deposits in CVB3-infected cardiomyocytes (30), suggesting that tissue deposition of iron in the heart may be a more prominent process during pathological inflammatory processes than previously appreciated.

CMR is the current gold standard for non-invasive detection of myocarditis and changes in T_1_ and T_2_ relaxation times are commonly used to detect parameters of inflammatory damage in the heart including edema, immune cell infiltration and fibrosis. Figure 5 illustrates how pathological changes in tissues affect CMR parameters measured as part of the Lake Louise criteria (LLC), which represent the first attempt to define a non-invasive diagnostic framework for myocarditis (31). The LLC protocol includes: (i) T_2_ weighted images to detect myocardial edema by measuring the myocardial T_2_ intensity change normalized to skeletal muscle (T_2_-STIR), (ii) early gadolinium enhancement (EGE) to detect reactive hyperemia, and (iii) late gadolinium enhancement (LGE) to assess tissue injury and/or fibrotic remodeling. In comparison, the approach taken in this study includes the measurement of potential hemorrhage/iron deposition, which strongly influences T_1_ and T_2_ and may complicate interpretation based on these parameters only. Yet, CMR-based detection of iron is performed routinely in patients with suspected cardiac iron overload (11) and β-thalassemia (32). A CMR imaging approach to detect myocardial iron in a mouse model of β-thalassemia has been reported recently (33).

**Figure 5:**
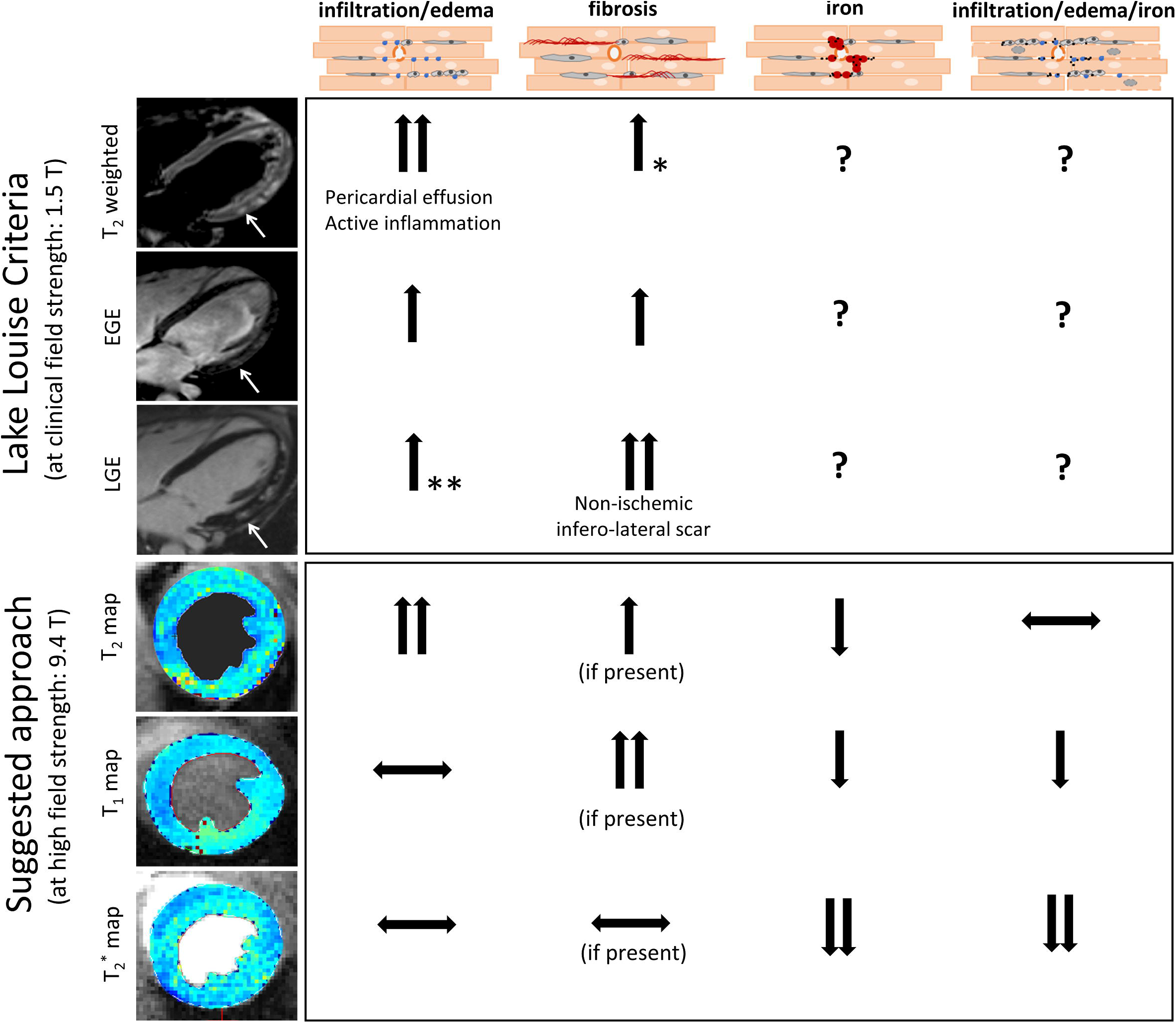
Comparison of the CMR protocol presented in this study to the established diagnostic approach of myocarditis in human patients based on Lake Louise Criteria (LLC). LLC-based measurements in human patients using clinical MRI field strengths. Edema leads to increased T2 weighted signal and increased T2 relaxation times suggestive of acute inflammation and or pericardial effusion. Hyperemia leads to increased early gadolinium enhancement (EGE) due leakage of gadolinium from capillaries. Necrosis or fibrosis can be identified by late gadolinium enhancement (LGE) as gadolinium will cross the damaged myocyte membrane (necrosis) or will accumulate in the extended extracellular space (fibrosis). Example CMR images illustrate an example of increased edema (increased T2), capillary leakage (increased EGE) and non-ischemic scar (positive LGE) at the infero-lateral wall of a human myocarditis patient. * Fibrosis can also non-specifically affect T2 values, however T2 alone is not sufficient to diagnose fibrosis but needs to be combined with T1 or LGE measurements. * * LGE may be enhanced in the presence of edema, however should not be used as a marker to identify myocardial edema, which should be done using T2 mapping. The approach proposed in this study and optimized on Resiquimod-treated mice with hemorrhagic myocarditis, provides a contrast-agent free CMR protocol able to detect inflammatory damage and allows correct interpretation of pathology despite potential interference from iron with the traditional T1 and T2 measures. In this model, edema increased, while iron decreased T2 relaxation times, and values appeared unchanged in the presence of both edema and iron due their opposing effects. Native T1 did not change in the presence of edema, and decreased in the presence of iron. It is expected to increase in the presence of fibrosis. T2* is strongly decreasing in the presence of iron and is not expected to change significantly in response to edema or fibrosis.

The mouse model used in this study has previously been described at chronic stage as a model of systemic autoimmunity (12). Yet, in the early acute phase it is likely to more closely mimic physiological responses to a viral infection due to the stimulation of the TLR-7 pathway. TLR-7 is a pattern recognition receptor involved in recognition of single-stranded RNA of viral origin thus crucial in host defense against viral infections (34). It is conceivable, that artificial over-activation of this virus defense system may cause the same phenomena as seen in severe complications of viral infection. This may include disseminated intravascular coagulation (DIC) due to acute systemic platelet activation leading to thrombocytopenia and internal hemorrhage (35). Notably, in an lymphocytic choriomeningitis virus (LCMV)-infection model, it was shown that only a severe drop in platelet count of more than 85% is necessary to cause hemorrhages (36) and even severely thrombocytopenic mice only develop local hemorrhages at sites of inflammation (37), which might explain the cardiac specificity of hemorrhages in Resiquimod-treated CFN mice. Systemic tissue damage incurred during this acute inflammatory phase, may then trigger the subsequent chronic autoimmune response observed in Resiquimod treated-mice. Both viral infections and tissue damage are known triggers of autoimmunity (38).

Using parametric CMR imaging, we demonstrate the feasibility to detect diffuse inflammatory infiltration and edema in mice with acute cardiac inflammation, and identify the presence of interstitial iron as a result of hemorrhage, which is paralleled by a significant reduction of T_1_ and T_2_ relaxation times. Based on the above, we have identified two main CMR phenotypes in hearts affected by inflammation and hemorrhage: 1) Increased T_2_ values with normal T_1_, T_2_^*^ representing infiltration/edema with normal levels of iron in the tissue suggestive of an acute ongoing inflammatory process as seen in classical myocarditis. 2) Low T_2_*, T_1_, T_2_ values representing infiltration/edema with increased levels of iron in the tissue suggestive of inflammatory changes including vascular damage, red blood cell extravasation and tissue iron deposition. This may need to be taken into consideration when designing myocarditis imaging protocols to avoid false negative results.

CMR imaging in mice is challenging due to limitations in achievable spatial resolution, fast heart motion, shortened relaxation times and strong field inhomogeneity coming with higher field strength (39). In addition, imaging procedures often rely on contrast agent application and lengthy acquisition protocols, limiting the opportunity of longitudinal measurements for paired analysis for welfare reasons. In a model of acute hemorrhagic viral infection, this is particularly important at acute disease stage, to avoid dehydration and excessive bleeding.

Notably, the relaxation times reported in this study will be different at the lower magnetic field strengths typically used in the clinic (i.e. 1.5T, 3T). Specifically, T2 relaxation times are longer, whereas T1 relaxation times are expected to shorten with decreasing magnetic field strength. T2* is influenced by the local magnetic susceptibility of interstitial iron and this effect is weaker with lower clinical field strength. This results in increased T2* values compared with preclinical CMR systems with higher magnetic field strengths. Based on this, clinical CMR systems may be more sensitivity to detecting edema (T2) and less sensitive to iron deposits measured by T2*, which may make the influence of iron on T2 measurements less pronounced.

In summary, the CMR protocol established in this study allows for detailed characterization of diffuse inflammatory damage in the mouse myocardium and controls for potential hemorrhage and iron deposition. Importantly, multi-parametric measurements of myocardial relaxation properties based on T1, T2 and T2* mapping is also a potential clinical strategy for the detection of diffuse immune-mediated damage, which also controls for the presence of hemorrhage.

## Author contributions

N.B. and S.S. conceived the study and designed experiments. N.B., S.S., A.P., I.S., A.L., R.C. J.B. and O.D. performed and analyzed experiments. N.B., S.S. and A.L. wrote the manuscript. S.E.H., N.R., L.Z. and S.S. provided financial support. S.E.H., L.Z., S.K.P, M.G.H and N.R. advised on project design and revised the manuscript.

---

**Table 1: Correlation of regional T_1_, T_2_ and T_2_^*^ relaxation times with their respective histological score.** Pearson’s Correlation was used to assess significance of correlation. * Correlation is significant at the 0.01 level (2-tailed). * * Correlation is significant at the 0.05 level (2-tailed)

**Supplementary Figure 1:** Improved multi-parametric approach to assess tissue damage by T_1_, T_2_ and T_2_^*^ mapping. Representative T_1_, T_2_^*^ and T_2_ maps acquired at base, midsection and apex of healthy mice showing the fesibility of this approach.

**Supplementary Figure 2:** MRI mapping parameters in the liver appear largely normal. Potential iron deposition and inflammatory damage to the liver in Resiquimod treated mice was assessed by T_1_, T_2_ and T_2_^*^ measurements in MRI.

