## Supplementary figures and images for "Detection of acute TLR-7 agonist-induced hemorrhagic myocarditis in mice by refined multi-parametric quantitative cardiac MRI"

### supplementary figure 1

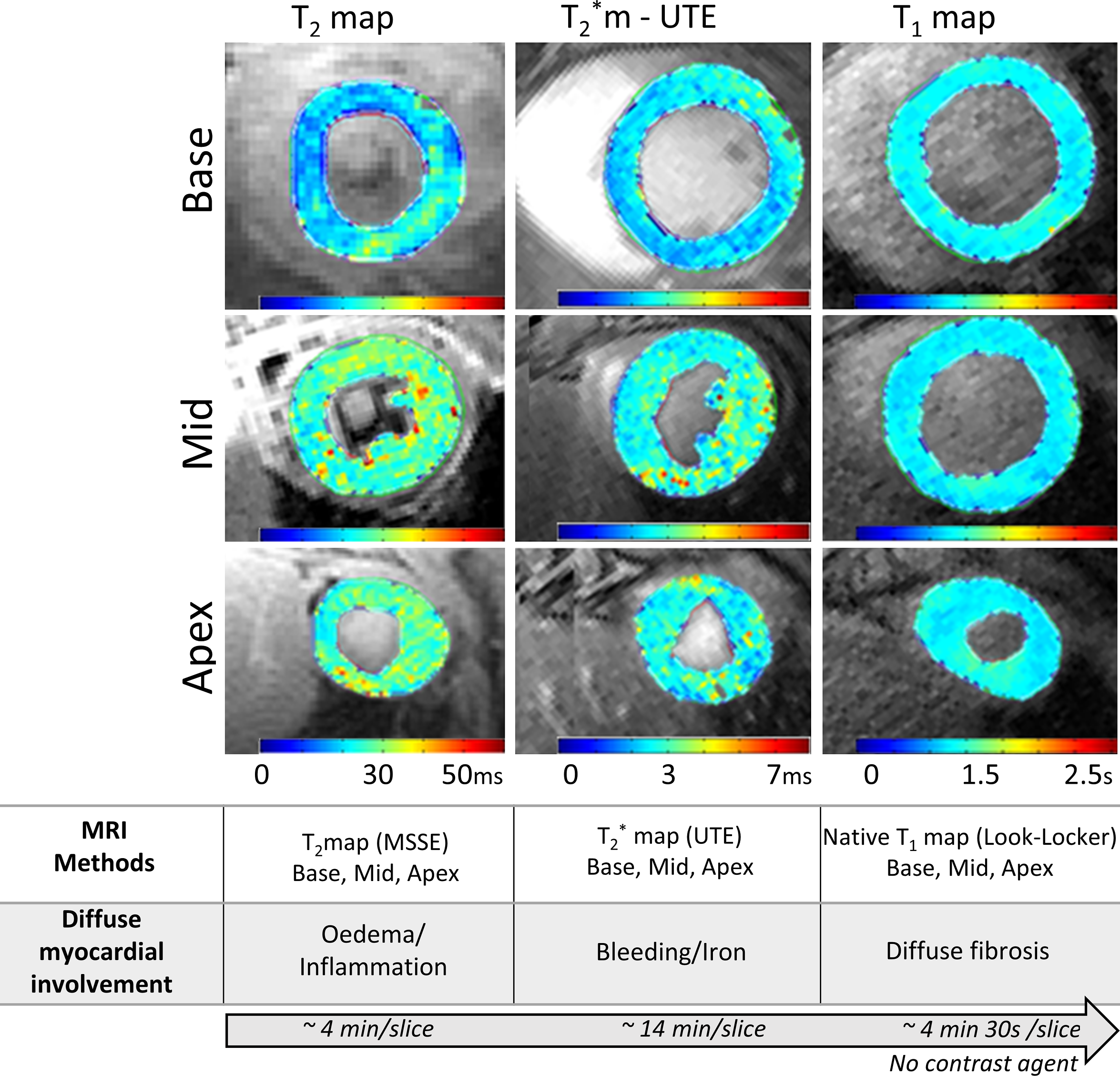

### supplementary figure 2

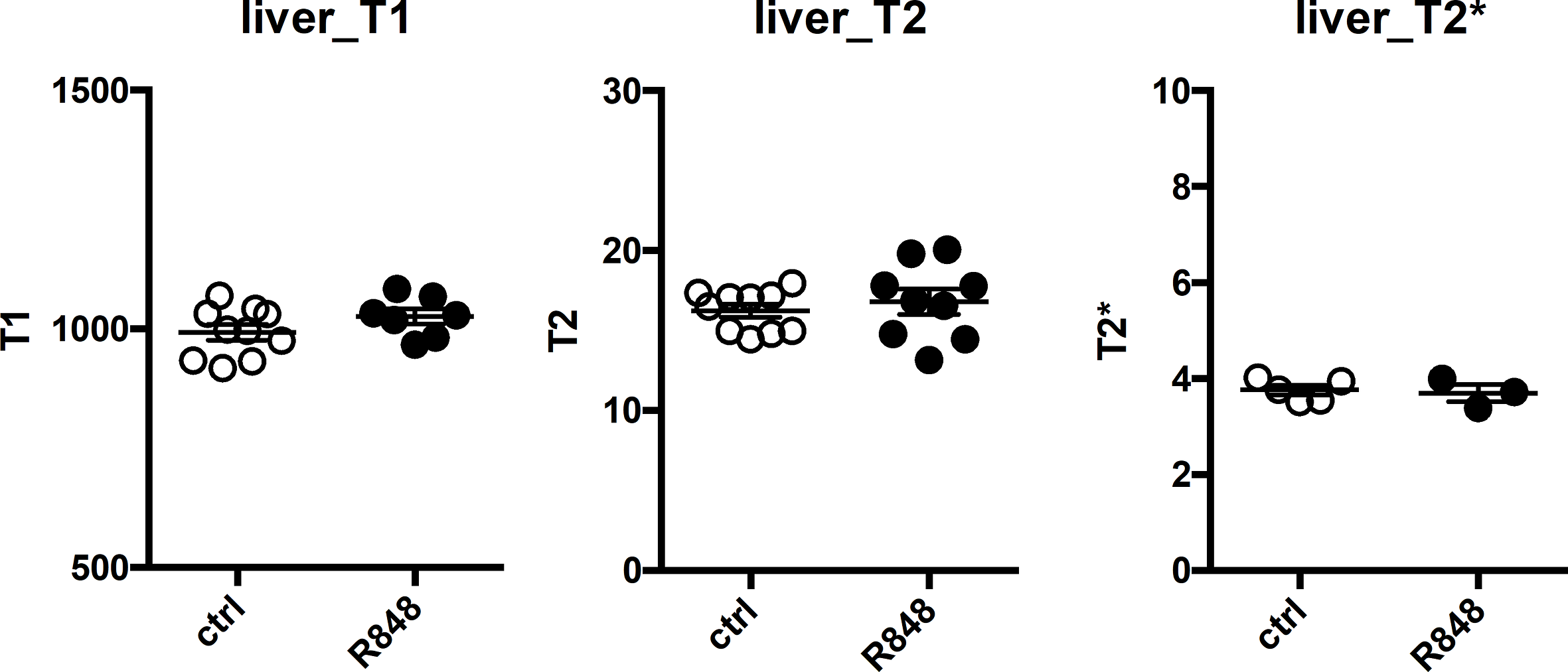
